# Topological inference from spontaneous activity structures in FMRI videos with peristence barcodes

**DOI:** 10.1101/809293

**Authors:** Arjuna P.H. Don, James F. Peters, Sheela Ramanna, Arturo Tozzi

## Abstract

Spatio-temporal brain activities with variable delay detectable in resting-state functional magnetic resonance imaging (rs-fMRI) give rise to highly reproducible structures, termed cortical lag threads, that can propagate from one brain region to another. Using a computational topology of data approach, we found that Betti numbers that are cycle counts and the areas of vortex cycles covering brain activation regions in triangulated rs-fMRI video frames make it possible to track persistent, recurring blood oxygen level dependent (BOLD) signals. Our findings have been codified and visualized in what are known as persistent barcodes. Importantly, a topology of data offers a practical approach in coping with and sidestepping massive noise in neuro data, such as unwanted dark (low intensity) regions in the neighbourhood of non-zero BOLD signals. A natural outcome of a topology of data approach is the tracking of persistent, non-trivial BOLD signals that appear intermittently in a sequence of rs-fMRI video frames. The end result of this tracking of changing lag structures is a persistent barcode, which is a pictograph that offers a convenient visual means of exhibiting, comparing and classifying brain activation patterns.

## Introduction

Point clouds are a natural outcome of a topology of data approach in tracking intermittent as well as persistent BOLD signals in different sections of the brain. A point cloud is a collection of sampled pinpointed places in a subregion [20]. A topology of data circumvents noise in data and focuses on those data that persist over time [15]. Computational topology of data approach provides a practical approach in isolating, measuring, and classifying persistent lag structures BOLD signals in each rs-fMRI video frame that are mapped to point clouds in a finite, bounded region in an *n*-dimensional Euclidean metric space. It has been observed that brain activity in one region of the brain propagates to others with variable temporal delay [25, 27, 33], giving rise to brain activity lag (delay) structures. Lag threads are temporal sequences of propagated brain activities [27]. Lag structures in fMRI video frames are rich source of point clouds. Triangulated brain point clouds are a source of brain activation area shapes that appear intermittently in different cortical regions during a rs-fMRI video such as the four videos provided by [27]. Selected barycenters of triangles in a triangulated cortical point cloud are connected to form vortexes covering each brain activation region with its own distinctive shape. Each vortex in a triangulated rs-fMRI video frame is a collection of connected cycles (see, *e.g.*, the lower half of Fig. 1) that make it possible to approximate, measure,track and compare brain activation region shapes. A vortical view of brain activity first appeared in W.J. Freeman [18]. Each vertex in a triangulated brain activation region point cloud is represented by a feature vector containing useful information such as activation region area and representative Betti number [22]. Traditionally, Betti numbers are either counts of surface holes or cycles. A particularly useful form of Betti number is a count of the number of connected vortex cycles covering an activation subregion [28]. Tracking the appearances of the Betti number of a brain activation vortex containing a particular number of connected cycles leads to the construction of a persistence barcode (see the top half of Fig. 1) in which a Betti number appears in a video frame, disappears afterward and possibly reappears one or more times in later video frames. 2D (planar) as well as 3D (volumetric) persistent barcodes provide an easy-to-read means of tracking intermittent BOLD signals in a sequence of rs-fMRI video frames.

**Figure 1.**
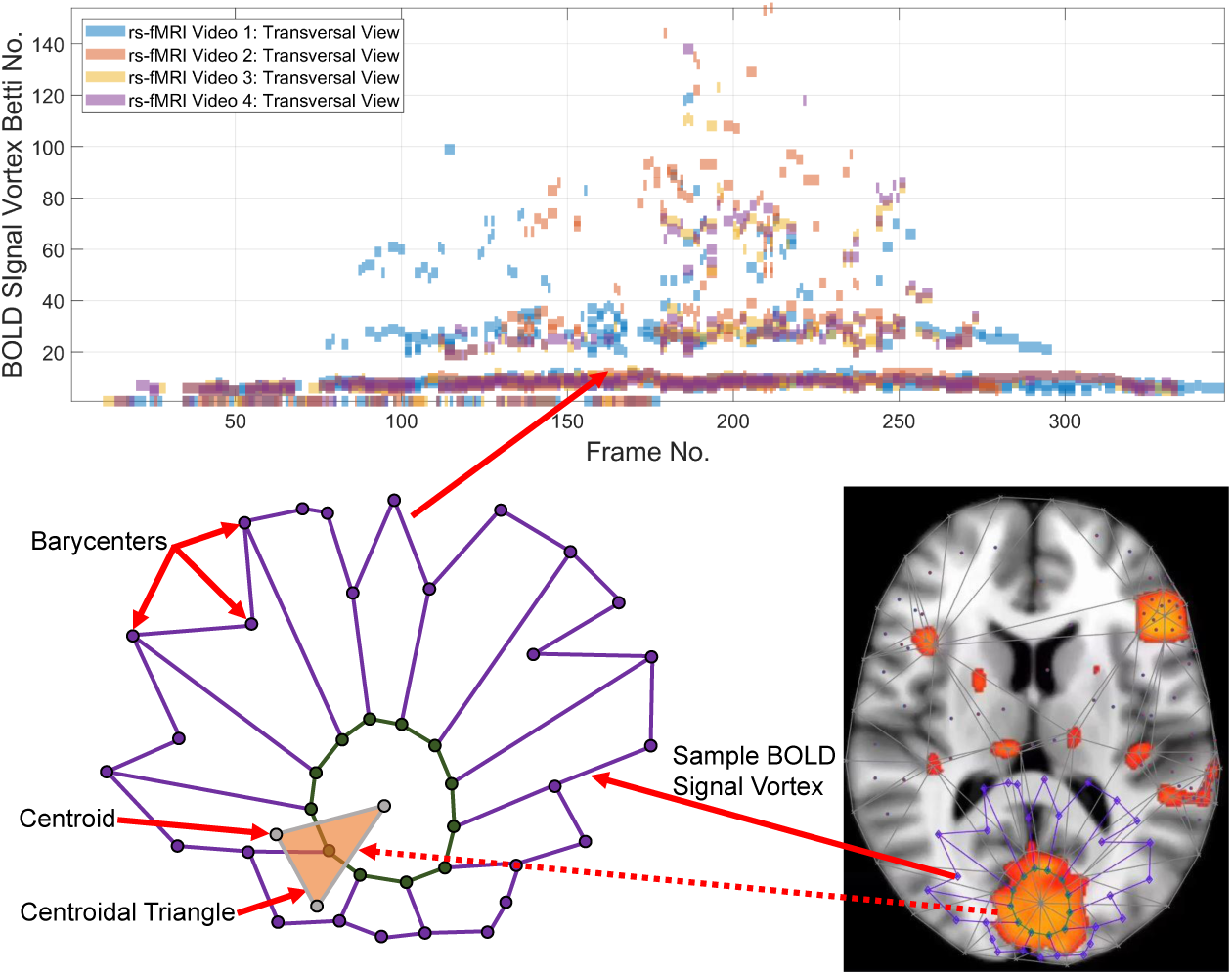
Betti number for a rs-fMRI BOLD signal vortex on a Transversal view of a brain region.

The origin of topological data analysis and persistent homology can be traced back to H. Edelsbrunner, D. Letscher and A. Zomorodian [16, 17]. A common approach is to build a continuous shape (graphs) on top of data to detect complex topological and underlying geometric structures [5, 7]. This shape is called a simplicial complex or a nested family of simplicial complexes and the process of shape construction is commonly referred to a filtration [35]. One of the fundamental tools in computational topology is persistent homology [34], that is a powerful tool to compute, study and encode efficiently multiscale topological features of nested families of simplicial complexes and topological spaces [14]. Earlier studies of brain networks primarily focused on a graph-theoretic approach where the brain regions and their connections are encoded as a graph (i.e., a network of nodes and edges) and cycles (representing complex behaviours). Such networks were modeled and analyzed with methods such as Q-modularity [26] or with network measures such as betweeness centrality [4]. Brain networks with weighted edges where problems of selecting thresholds for edge weights and dealing with sparse edges can be found in [1, 32]. This has led to application of persistent homology to the problem of determining multiple thresholds derived from more than one network. A brain network can be considered as the 1-skeleton of a simplicial complex, where the 0-dimensional hole is the connected component, and the 1-dimensional hole is a cycle [9]. The number of k-dimensional holes of a simplicial complex is its k-th Betti number. Persistent homology-based multiscale hierarchical modeling was proposed in [9, 22, 24, 30, 31] to name a few. Here, graph filtration is used to build these networks in a hierarchically manner. Filtration is the process of connecting edges to form a graph.

Betti numbers computed during this filtration process have been used for further statistical analysis such as Pearson correlation to MRI image data [8] and various metrics for similarity and distances assessments [9, 23]. 0-dimensional holes (*B*_0_ or zeroth Betti numbers) have been computed during the graph filtration process and a persistent barcode has been constructed for subsequent statistical analysis [6, 10]. However, the *cycle* concept is extremely important to the study of behaviour diffusion and integration of the brain network. In persistent homology, cycles are measured using the first or one-dimensional Betti Number (*B*_1_). In [9], the one dimensional Betti numbers are used to measure cycle and the significance of the number of cycles is evaluated using the Kolmogorov-Smirnov (KS) distance.

In contrast to earlier approaches, we use computational geometry to detect lag thread shapes in fMRI video frames using the first Betti number to measure connected cycles called vortexes derived from the triangulation of brain activation regions. We build on our previous work in [13, 28, 29] to evaluate real BOLD resting state rs-fMRI videos from [27]. Here, the video frames are processed directly to obtain the first Betti numbers by triangulating the transversal, sagittal and coronal sections of the fRMI video frames and constructing vortexes through a process of filtration. Rather than constructing graphs of brain networks and analysing cycles in the network, we analyze cycles by processing the video frames. The vortexes correspond to the changing activation areas in the video frames. The number of vortexes in a frame represents the most relevant areas of change. Higher betti number values imply that the change is closer to the center of the section of the brain. To the best of our knowledge, this method of detecting cycles in persistent homology is novel.

## Theory

The basic approach is to introduce a free Abelian group representation of intersecting filled polygons on the barycenters of the triangles of vortex nerves [13, 28]. Each vortex is a collection of connected cycles called a vortex nerve. Of particular interest are those nerves that have the maximal collection of triangles of a common vertex in the triangulation of a finite, bounded planar region. In our case, the planar region is a video frame. A vortex nerve results from the triangulation of the sections of the fMRI brain video. A centroid (also called a seed point), is used as a vertex in the triangulation of video frame image. A barycenter on a such a triangle is in an image region between the dark regions which we refer to as *holes*. The concept of a hole is crucial to this work since edges and interior each vortex nerve reveal paths of reflected light between dark regions (holes). As a result, connected barycenters model paths for light from either reflected or refracted light from visual scene surface shapes recorded in a video frame. A one-dimensional Betti number is a count of the number of generators in a finite free Abelian group [21, p. 151].

In our case, a Betti number tells us the number of generators in the frame vortex nerve. A repetition of the same one-dimensional Betti number across a sequence of consecutive frames informs us that a similar shape recurs in these frames. The Betti number of a vortex containing 2 main cycles connected by *k* edges attached between the pair of vortex cycles equals k + 2 [13]. For example, the Betti number of the vortex in the lower half of Fig. 1 equals 14, since this vortex contains two main overlapping cycles of 12 single edge cycles attached between the main cycles. A vortex containing *n* ≥ 2 connected main cycles and a total of *m* connecting edges yields a Betti number equal to *m* + *n*. Each one-dimensional Betti number in a persistent barcode [19] represents the number of connected cycles covering an activation area of the brain. Recall that a one-dimensional Betti number is a count of the number of connected cycles in a graph. In our case, for example, a single edge attached between a pair of vortex cycles is modelled as a simple bi-directional cycle. This topology of data pictographs is useful in representing the persistence of the brain activation region shapes found in rs-fMRI brain video sections. The focus here is on the persistent Betti numbers across sequences of triangulated video frames. Each Betti number is mapped to an entry in a persistent barcode (see top half of Fig. 1).

## Materials and methods

This section briefly introduces the methods used to derive a vortex cycles on triangulated video frames (steps 0 through 6) and their Betti numbers (step 7), which are used to construct persistent barcode (step 8) for rs-fMRI videos. The fMRI videos (of 688 subjects) used in this work were obtained from the Harvard-MGH Brain Genomics Superstruct Project [3]. Each video contains 360 frames that exhibit the propagation of BOLD signals in the sagittal, transversal and coronal sections of the human brain (see, *e.g.*, the middle row of Fig. 2). Let *K* be a rs-fMRI video frame. The steps to obtain triangulation vortexes covering the brain activation regions shown in Fig. 2 are as follows: **0**^**0**^ Find the centroids (centers of mass) of the background regions (in mathematical morphology, such regions are termed holes) surrounded by foreground vortexes in brain activation regions. Holes are identified by binarizing each video frame. In a rs-fMRI video frame, the holes are in background regions containing dark (low intensity) voxels and the foreground regions are filled with mainly orange voxels. **1**^**0**^: *Triangulate* on the centroids of the tiny dark regions (holes) inside the brain activation regions in *K*. A sample centroidal triangle is shown in Fig. 1. **2**^**0**^: Find the *barycenter* of every centroidal triangle in Δ *K*. Each barycenter is a voxel representing a non-low BOLD signal situated between centroids (low intensity voxels). **3**^**0**^: Connect the *barycenters* where there is the greatest concentration of centroidal triangles Δs with a common vertex. Recall that the vertex common to a collection of triangles is an example of an Alexandroff nerve [2] (see, *e.g.*, the collection of centroidal triangles with a common vertex covered by the inner vortex cycle in Fig 1). In this work, the focus is on finding maximal Alexandroff nerves. The set of barycenters surrounding the centroid of an Alexandroff nerve forms a *cycle C*_0_. Notice that cycle *C*_0_ will always be in the interior of a brain activation region containing high intensity (orange) voxels. **4**^**0**^: Connect the *barycenters* of centroidal triangles {Δ} along the *boundary* of *C*_0_. This collection of circularly connected edges constitutes a new cycle called *C*_1_. Cycle *C*_1_ usually overlaps the boundary of a high intensity brain activation region. **5**^**0**^.**1**: Connect each barycenter in *C*_0_ to a neighbouring barycenter on *C*_1_. **5**^**0**^.**2**: Repeat steps 4.0 and 5.0.1 for each cycle constructed on the barycenters outside the boundary of *C*_1_ and inside the boundaries between subregions containing high BOLD signals. **6**^**0**^: Repeat steps 1 through 5 until there are no more vortexes covering subregions containing nonzero BOLD signals. The end result is a collection of connected nesting non-concentric cycles that form a *vortex* (see, *e.g.*, the sample 14-cycle vortex in Fig. 1, *i.e.*, 2 large connected cycles *C*_0_ and *C*_1_ attached to 12 signal edge cycles). Once these vortexes are generated, the next step is to compute the one-dimensional Betti number for each vortex that has been found in each of the triangulated video frames. Notice that there is usually more than one vortex in video frame. **7**^**0**^: Count the number of non-single edge (main) cycles in a vortex plus the number of signal edge cycles connected between the main vortex cycles to obtain a vortex Betti number. This process is repeated for each brain section in every frame containing sagittal, trasversal and coronal brain sections in each of the sample rs-fMRI videos. **8**^**0**^: To construct a Betti number-based persistent barcode, insert a bar in a pictograph (an easy-to-read visualization of brain activity instants accumulated in what is known as a homology barcode [20]), using rs-fMRI video frame number (x-axis) and its corresponding Betti Number (y-axis) (shown in the top half of Fig. 1).

**Figure 2.**
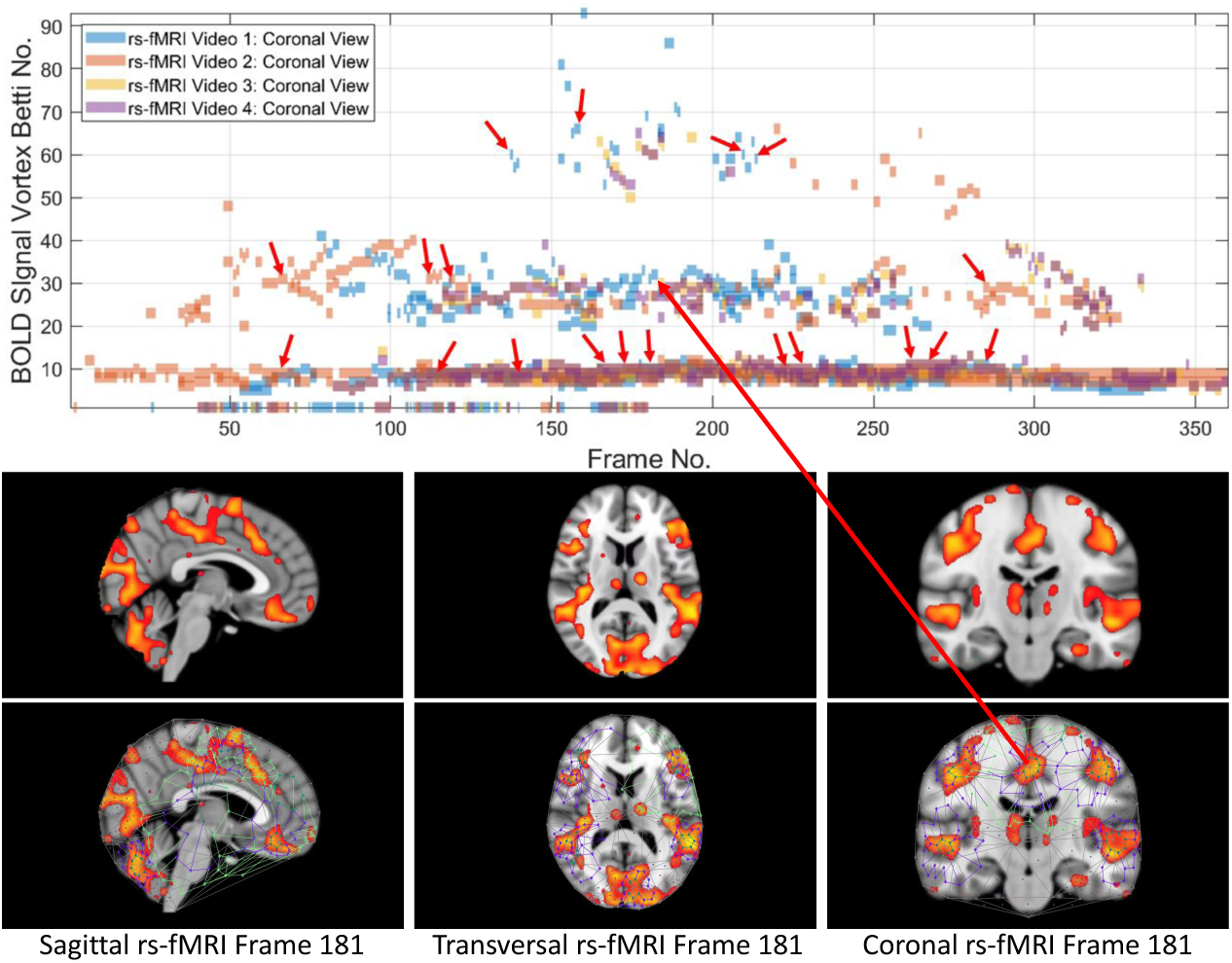
Coronal persistence barcode with Betti numbers as vortex cycle counts in a rs-fMRI video frame (See text for further details)

The gaps between the sequences of contiguous bars are important, since gaps in a particular row of pictograph bars indicates rs-fMRI video frames that do not have the same level of brain activity represented by bars in the row. The proximity of the bars (not necessarily contiguous) in a pictograph row call attention of BOLD signals that are close to each other, temporally. Repeated bars in a pictograph row indicate a repeated (persistent) level of brain activity recorded in the rs-fMRI video frames. An examples of a pictograph row containing multiple, contiguous bars can be seen in row 30 (Betti numbers = 30) in Fig. 2. Contiguous bars in a pictograph row indicate the closeness in time of the corresponding brain activity. Examples of pictograph rows containing multiple, non-contiguous bars can be seen in rows 10 and 60 (Betti numbers = 10 and in Fig. 2. A byproduct of the inspection of a sequence of contiguous bars (Betti numbers) in a persistent barcode row can lead to the production of a reduced rs-fMRI video containing only video frames with either activation sub-regions with the same Betti number or a new video containing frames with activation sub-regions, each with a different Betti number.

## Results

### Part 1. Derivation of Persistent Brain Activation Subregion Signature

Each triangulated BOLD signal propagation subregion has a signature defined by the vector (frame number, Betti number, inner vortex cycle area). This leads to the production of four triangulated rs-fMRI videos available at [12] that are part of the University of Manitoba Vortex Signature Project. This triangulation provides a computational geometry perspective on the structure of brain activation subregions. Vortexes have been derived from each of the triangulation of the BOLD signal activation regions in each of the brain sections in the video frames in the four rs-fMRI videos from [3]. A straightforward outcome of the derived vortexes is a rich source of new means of describing individual BOLD signal activation regions as well as a bridge to various forms of descriptions of lag thread. For example, each vortex has a Betti number (count of the number of connected cycles) and various cycle areas. Each of the three brain regions in each frame in the Harvard Brain Genomics rs-fMRI videos has its own vortex and, consequently, its own Betti number. Typically, each video frame will have more than one Betti number derived from vortexes on the multiple brain activation subregions. Betti numbers provide a computational topology perspective on the structure of brain activation subregions. In this study, the focus is on the area of the inner cycle of a BOLD signal subregion vortex. This is the case, since each inner cycle lies entirely within the interior of an activation subregion. Hence, an inner cycle area is a reliable approximation of the brain activation area. In sum, Betti numbers and inner vortex cycle area help gauge the extent of an activation subregion. Considered either separately or taken together, a vortex on brain activation subregion provides the basis for a subregion signature, *i.e.*, a distinctive characteristic of a brain activation subregion in a rs-fMRI video frame. For example, (frame number, Betti number) = (140, 60), (200, 60), (210, 60) provide a signature for the coronal brain region in frames 140, 200 and 210 in Fig. 2. Betti number 60 is an example of a brain activation subregion characteristic that persists over a sequence of video frames. The vector (frame number, Betti number, inner vortex cycle area) = (180,40,10 mm^2^)depicts a brain activation subregion in the sagittal brain region in Fig. 3.

**Figure 3.**
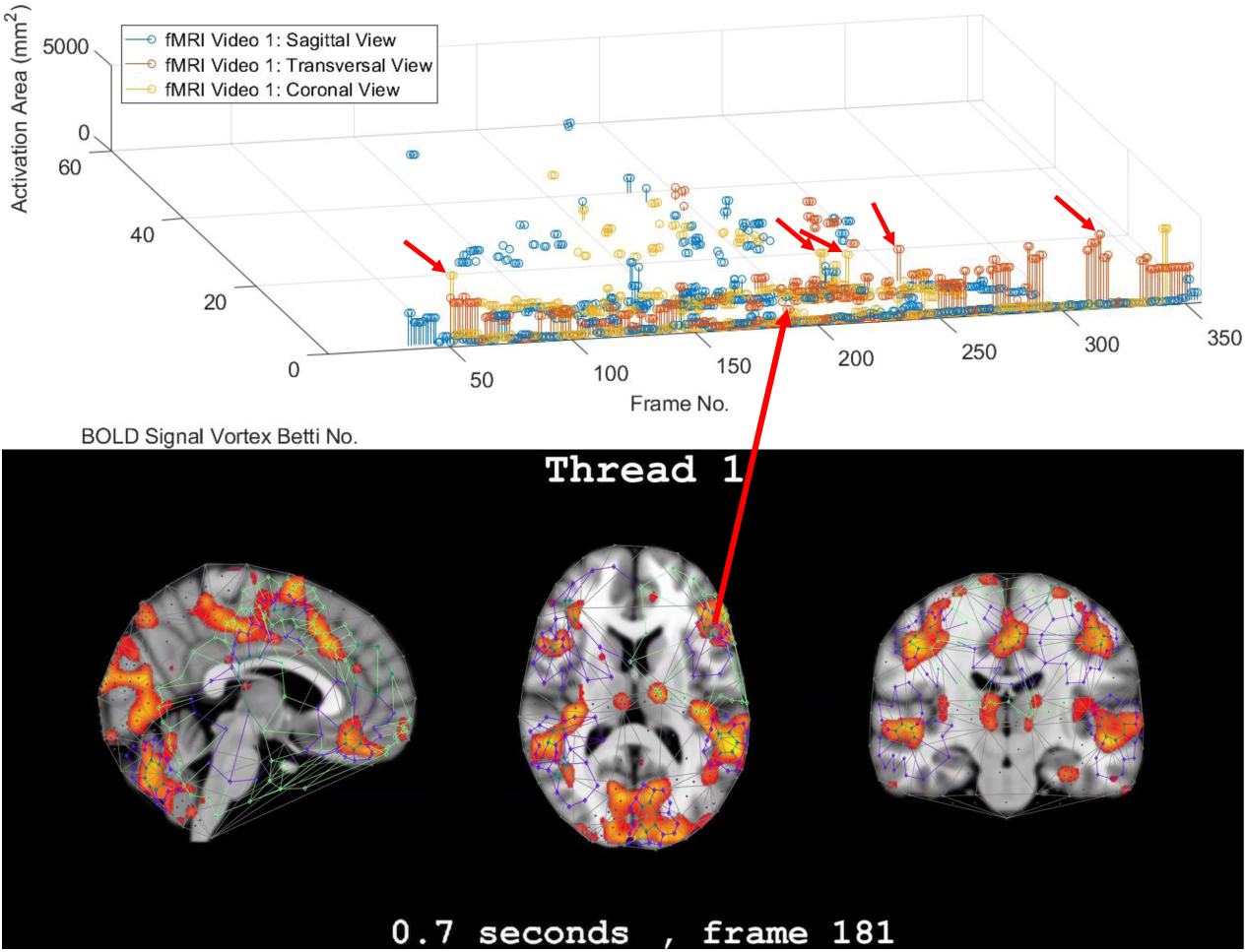
Frame-Betti number-Area for a rs-fMRI BOLD signal vortexes on 3 brain regions.

### Part 2. Betti number-based rs-fMRI lag thread pattern

A repetition of the same Betti number for the same brain region across multiple video frames defines a lag thread pattern. Let ℬ_*s*_, ℬ_*t*_, ℬ_*c*_ be Betti numbers for the sagittal, transversal and coronal brain regions. For example, ℬ_*s*_ = ℬ_*t*_ = ℬ_*c*_ = 100 defines a lag thread pattern for multiple frames in Fig. 3.

### Part 3. Confirmation of highly reproducible lag thread topography

From 3D barcodes in Fig. 3 as well as in Fig. 4, Betti number-area patterns can be detected within frames in the same video. That is, one can find many examples of brain activation subregion Betti number (and corresponding subregion area in a lag thread in one video frame) that are reproduced in a lag thread in a different video frame. In other words, the Betti number-area combination persists across different frames. Many examples of this Betti number-area persistence phenomenon can be detected in the two sample 3D homology barcodes when compared with similar 3D homology barcodes derived from the frames in the videos available at the University of Manitoba Vortex Project at [12]. These persistent repetitions in the topography detected in different lag threads confirm the observation that *there are commonalities in signal propagation within each lag thread* [27].

**Figure 4.**
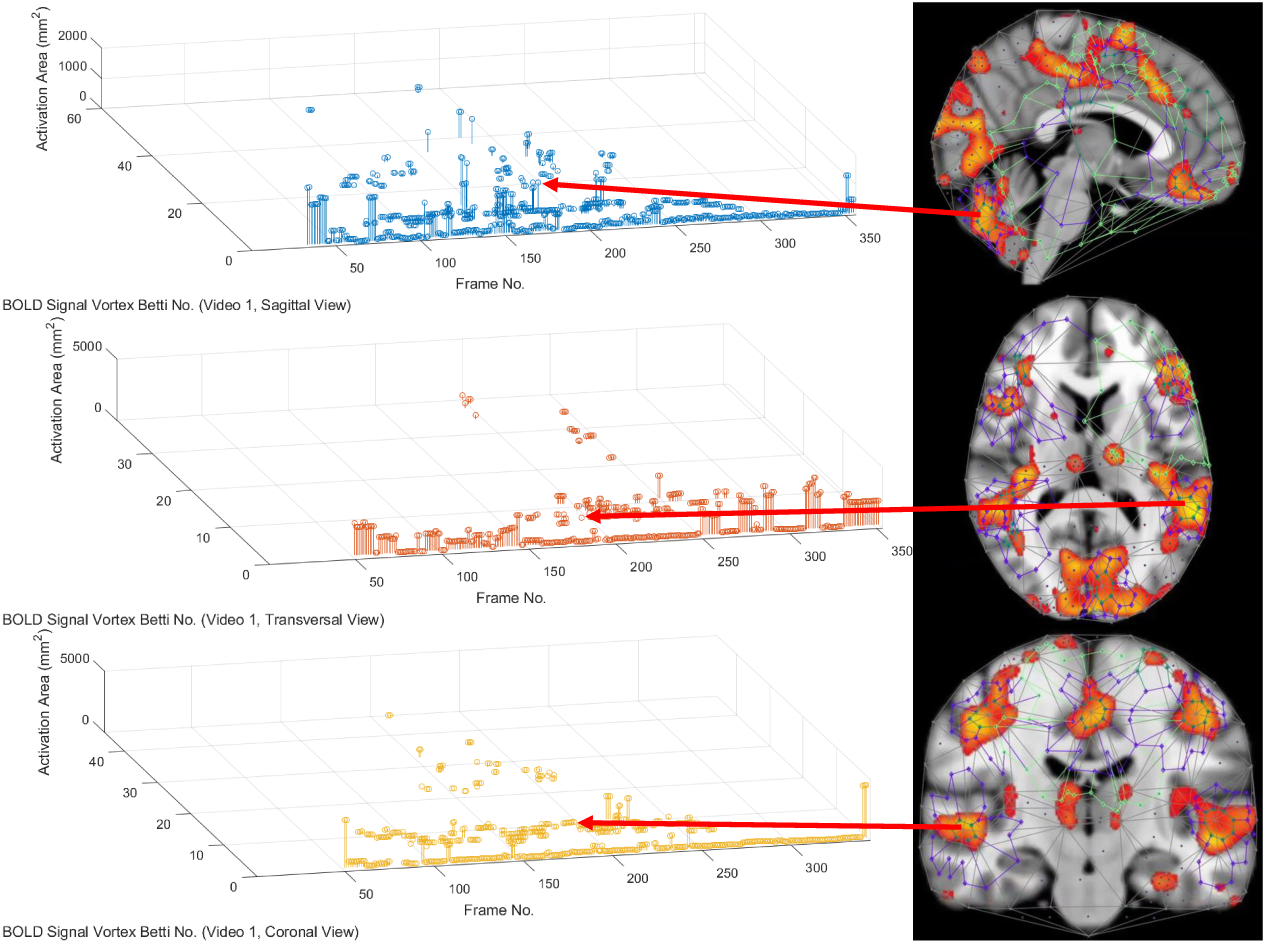
3D persistence barcode for 3 rs-fMRI BOLD signal propagated in brain regions.

### Part 4. rs-fMRI lag threads having descriptive proximity

A pair of objects are descriptively proximal (near each other), provided the objects have the same description [11]. A feature vector provides a description of an object. In this work, (Betti number, area) describes a subregion of a rs-fMRI brain region. Rather than a purely theoretical, abstract approach to descriptive proximity spaces, the focus here is on computational descriptively proximities. Briefly, computational descriptive proximity includes algorithms as well as structures such as set intersection, union and closure and proximity space axioms introduced in [28]. Descriptive proximites provide mathematical framework useful in measuring, comparing, and classifying (1) lag structures and threads across frames in the same video or (2) lag structures and threads across frames in different rs-fMRI videos. For example, in terms of (1), (ℬ_*t*_, inner vortex cycle area) = (100, 100 mm^2^) describes a brain activation subregion in the transversal brain section in frames 10 and 75 in Fig. 3. In terms of (2), the feature vector (ℬ_*s*_, ℬ_*t*_, ℬ_*c*_) can be used to describe and compare lag threads in frames across different videos. Again, for example, the feature vector (ℬ_*t*_, *area*) can be used to compare brain activation subregion areas with the same Betti number either across frames in the same video or frames in different rs-fMRI videos. This is an important advantage that accrues from the application of computational descriptive proximities.

## Discussion

### Summary of Findings

This study of the Betti numbers and inner vortex areas of rs-fMRI BOLD signal propagation subregions of the brain confirms and supplements earlier findings given in [27]. A main result of this study of the persistence of brain activation subregion features confirms the contention that the topography of lag threads is highly reproducible. Starting with the Betti number of connected cycles derived from triangulated brain activation regions found in rs-fMRI videos, it is apparent that the vortexes on brain activation subregions appear over and over in the lag threads across different rs-fMRI video frames. In other words, we find that there are commonalities in BOLD signal propagation within each lag thread.

The question whether intrinsic brain activity contains reproducible temporal sequences is revisited. It is confirmed that a human resting state fMRI (rs-fMRI) contains persistent (repeatable) highly reproducible lag structure. The answer to this question is given twofold. This is done first in terms of Betti numbers that are counts of the number of connected cycles in vortexes on triangulated brain activation subregions. We introduce a 2D persistence pictograph (barcode) that exhibits the appearance, disappearance, and repeated reappearance of Betti numbers across lag threads in sequences of rs-fMRI video frames. In addition, the reproducibility question is also answered in terms of the introduction of a video frame-Betti number-vortex cycle area combination in 3D persistence barcodes that facilitates a check on how often these features of lag threads appear during a rs-fMRI session.

### Concluding Remarks

This study considers Betti numbers that are counts of the number of connected cycles in a vortex on brain activation subregion. Another Betti number of interest and not considered here is the count of the number of holes (dark surface regions) in lag structures. In terms of the area occupied by a vortex on a brain activation subregion, we have only considered the area of the interior of the innermost of each vortex. Also of interest and of considerable importance is the area in the interior of other cycles that includes the inner vortex cycle. Future work would expand the derivation of persistent barcodes to include the zeroth as well as the oneth Betti numbers. In working toward the approximation of the area of a brain activation subregion shape, the areas in the interior of the other cycles (summing on the innermost vortex cycle area) would be considered.

## Acknowledgments

This research has been supported by the Natural Sciences & Engineering Research Council of Canada (NSERC) discovery grants 185986 and 194376, Instituto Nazionale di Alta Matematica (INdAM) Francesco Severi, Gruppo Nazionale per le Strutture Algebriche, Geometriche e Loro Applicazioni grant 9 920160 000362, n.prot U 2016 / 000036

